# Inconsistent estimates of hybridization frequency in newts revealed by SNPs and microsatellites

**DOI:** 10.1101/2023.01.21.525005

**Authors:** Aurélien Miralles, Jean Secondi, Maciej Pabijan, Wiesław Babik, Christophe Lemaire, Pierre-André Crochet

**Affiliations:** Zoological Institute, Braunschweig University of Technology, Mendelssohnstr. 4, Braunschweig, 38106, Germany; Institut de Systématique, Evolution, Biodiversité, (UMR 7205 Muséum National d’Histoire Naturelle, CNRS UPMC EPHE, Sorbonne Universités), CP30, 25 Rue Cuvier, 75005, Paris, France; Univ Lyon, Université Claude Bernard Lyon 1, CNRS, ENTPE, UMR 5023 LEHNA, F-69622, Villeurbanne, France; Faculté des Sciences, Université d’Angers, 2 Bd de Lavoisier, 49000 Angers, France; Institute of Zoology and Biomedical Research, Faculty of Biology, Jagiellonian University, Gronostajowa 9, 30-387 Kraków, Poland; Institute of Environmental Sciences, Faculty of Biology, Jagiellonian University, Gronostajowa 7, 30-387 Kraków, Poland; Université d’Angers, Institut Agronomique Rennes-Angers, INRAE, IRHS, SFR, QUASAV, 49000 Angers, France; CEFE, CNRS, Univ Montpellier, EPHE, IRD, Montpellier, France

**Keywords:** Broad sympatry, introgression, *Lissotriton vulgaris*, *Lissotriton helveticus*, SEQUENOME

## Abstract

Hybridization between the European smooth and palmate newts has recurrently been mentioned in the literature. The only two studies that attempted to quantify the frequency of hybridization and gene admixture between these two species came to strikingly opposite conclusions. According to Arntzen et al (1998, 42 allozymes), hybrids are rare in nature and introgression negligible, while according to Johanet et al (2011, 6 microsatellites), introgressive hybridization is significant and widespread across the shared distribution range. To clarify this question, we implemented high-throughput SNP genotyping with diagnostic biallelic SNPs on 965 specimens sampled across Europe. Our results are in line with Arntzen et al, since only two F1 hybrids were identified in two distinct French localities, and no further hybrid generations or backcrosses. Moreover, reanalysis of 78 of the samples previously studied by Johanet et al. (2011) using our SNPs panel could not reproduce their results, suggesting that microsatellite-based inference overestimated the hybridization frequency between these two species. Since we did not detect methodological issues with the analyses of Johanet et al., our results suggest that SNP approaches outperform microsatellite-based assessments of hybridization frequency, and that conclusions previously published on this topic with a small number of microsatellite loci should be taken with caution, and ideally be repeated with an increased genomic coverage.

## Introduction

Introgressive hybridization is a process of major interest for evolutionary biology. By blurring the lines between species, interspecific gene flow reveals the gradual nature of reproductive isolation and its possible reversibility, complicates species delimitation, and more generally calls into question the centuries-old categorical conception we may have of species (Taylor et al. 2006, Harrison & Larson 2014, Barraclough & Humphreys 2015, Moran et al. 2021, Kim et al. 2022). The transfer, *via* hybridization and backcrossing, of novel alleles from one species into the gene pool of another species also constitutes a non-negligible source of genetic diversity, which may positively or negatively impact the adaptive capacities of species (Barton 2001, Pfenning et al. 2016, Seabra et al. 2019, Steensels et al. 2021, Wacker et al. 2021). Importantly, anthropogenic hybridization is increasingly common and its impact on conservation issues has been the topic of much debate (Chan et al. 2019, Mc Farelane & Pembertone 2019, Hirashiki et al. 2021).

Hybridization in animals has for a long time been considered as an anecdotal phenomenon, but in recent years, a growing body of evidence has demonstrated that a broad range of animal species experience it during their history (Mayr 1963, Mallet 2005, Taylor & Larson 2019, Adavoudi & Pilot 2022). In amphibians, introgressive hybridization has been documented in numerous anurans (e.g. Dufresnes et al. 2021), possibly as a result of their frequently external fertilization reducing the efficiency of pre-zygotic reproductive barriers. In Eurasian Urodela (with internal fertilization), several well documented examples have been reported in the genera *Triturus* (Jehle et al. 2001, 2009, 2021, Cogălniceanu et al. 2020) and *Lissotriton*, where *L. vulgaris* and *L. montandoni* exhibit some dramatic genomic consequences of past introgression (Babik et al. 2003, 2005, Babik & Rafiński 2004, Gherghel *et al*. 2012, Zieliński et al 2013, 2014, 2016, Pabijan *et al*. 2017, Niedzicka *et al* 2017, 2020, Dudek *et al*. 2019). Prior to these works, several sources (Griffiths 1987, Arntzen et al 1998, Beebee et al. 1999, Schlüpmann et 1999) have also mentioned hybridization between the smooth and palmate newts, *Lissotriton vulgaris* and *L. helveticus*, two distantly related and non-sister species within the genus that diverged from each other relatively early (the divergence is imprecisely estimated between ∼15 Mya and 24 Mya (Rage & Bailon 2005, Steinfartz et al. 2007, Böhme 2010, Arntzen et al 2015)). The first work aiming at quantifying the extent of hybridization between these two species was based on multivariate analyses of 16 morphological traits and electrophoretic analyses of 42 protein loci (Arntzen et al. 1998). Although this work irrefutably demonstrated natural hybridization between *L. vulgaris* and *L. helveticus*, it concluded that this phenomenon was rare (only one F1 hybrid identified from a large sample (>5,000) of larvae, recently metamorphosed newts and adults) and no introgression was detected between the species (cf. Fig. 3 of Arntzen et al.). More recently, and in striking contrast to this first work, Johanet et al. (2011) used mitochondrial and microsatellite markers on ∼1,300 individuals from 37 sites across Europe and concluded that introgression was instead relatively widespread in the area of sympatry, with a frequency of hybridization of 1.7% and significant levels of introgression detected at most sites (73%) shared by both species. To determine the extent of hybridization and introgression between *L. vulgaris* and *L. helveticus* and to elucidate the causes of the discrepancy between the former studies, we reinvestigated the frequency of hybridization between the two species, using extended geographical sampling and, for the first time, high-throughput SNP genotyping.

## Material and methods

### Sampling

Currently, *Lissotriton vulgaris* has a broad Eurasian distribution, whereas *L. helveticus* is restricted to the western part of Europe (Wielstra et al. 2018; Sillero et al., 2014). Both species occur sympatrically across a wide area including Great Britain, the north of France, Switzerland, the Benelux countries and the west of Germany (Fig 1). Designed to take into account their respective distribution, our sampling includes a total of 965 individuals from 59 localities (=ponds): 29 samples from 11 allopatric localities for *L. helveticus* (in Spain and southern part France), 23 samples from 8 allopatric localities for *L. vulgaris* (in Norway, Sweden, Poland, Romania and Hungary), and 913 samples of both species from 40 sympatric localities (in Great Britain and northern France), of which 829 are from 32 localities where both species co-occur in syntopy (see Fig. 1 and Table 1). All these samples were collected during the reproductive phase (aquatic phase) and therefore correspond to viable adult individuals. Notably, our survey also includes a selection of 78 syntopic individuals that were previously analyzed using microsatellites by Johanet et al. (2011), including those that showed the most substantial rates of introgression.

**Fig 1.**
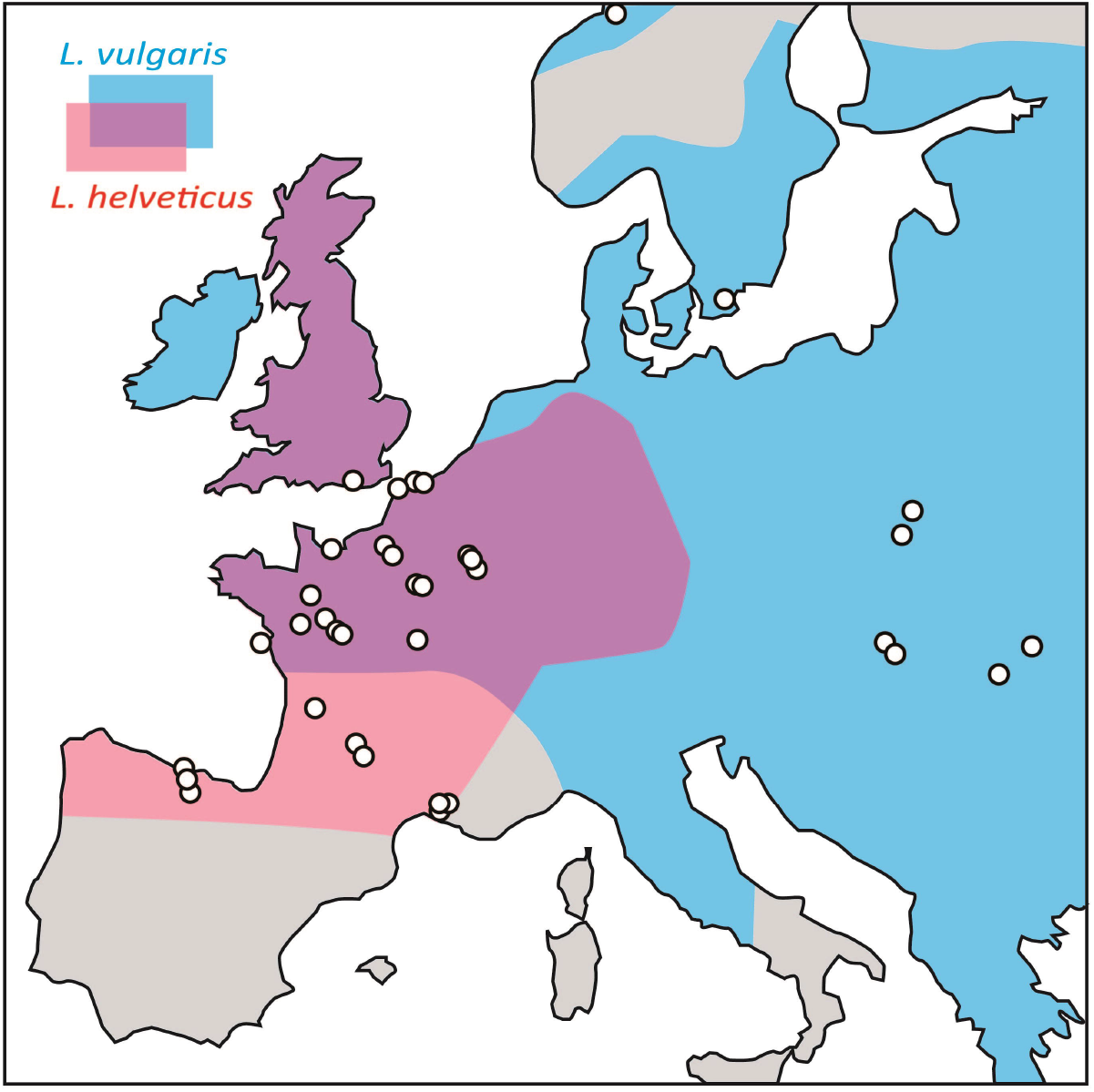
Population sampling. The red and blue areas indicate the approximate distribution of *Lissotriton helveticus* and *L. vulgaris*, respectively (sympatric distribution range represented in purple). Each white dot represents one or several sampled ponds from the same area (cf. Table 1 for details).

**Table 1.**
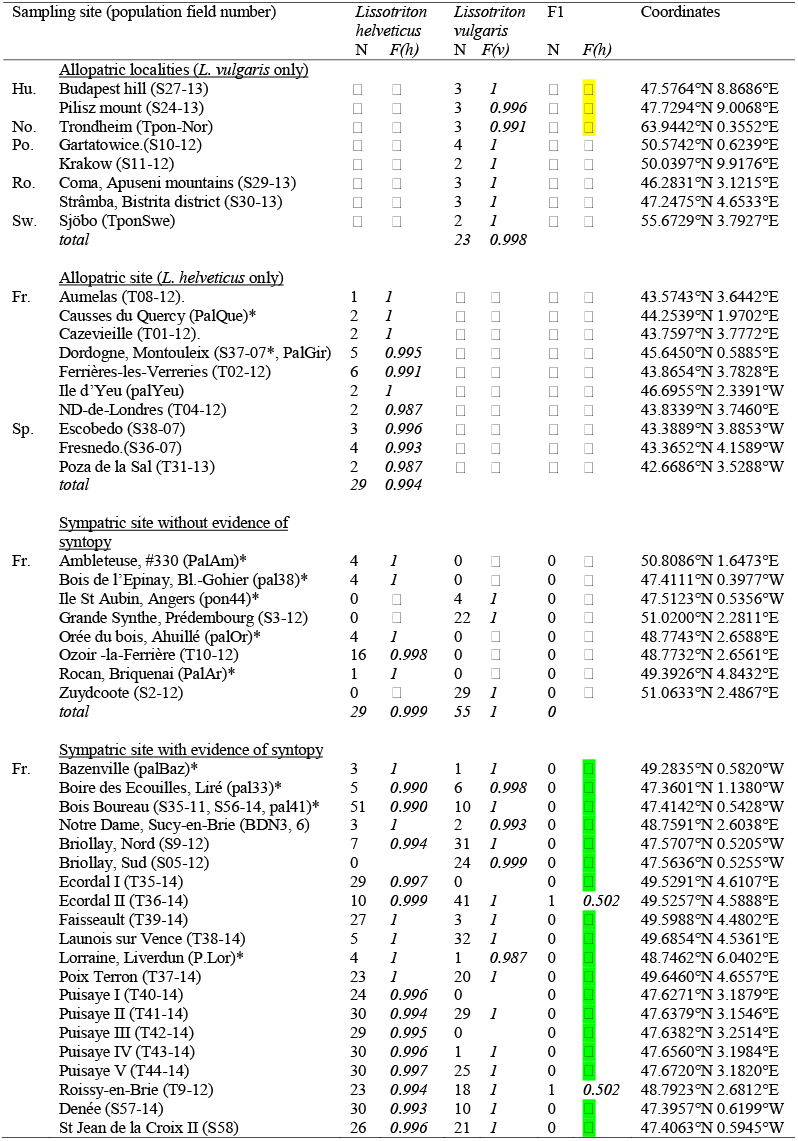
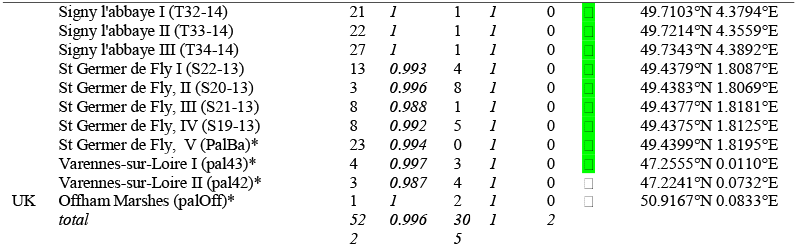
Geographic sampling of *Lissotriton helveticus* and *L. vulgaris*. For each locality, the number of samples (N, assignment to a given species based on the present results) and the overall frequencies of the diagnostic alleles for each species (*F(h)* and *F(v)*, respectively) are indicated (only *F(h)* frequencies are indicated for the two hybrid specimens). Abbreviations : Hu. Hungary, No. Norway, Po. Poland, Ro, Romania, Sw. Sweden, Fr, France, Sp. Spain, UK. United Kingdom ; F1 represents putative F1 hybrids. Localities including samples already analysed by Johanet et al (2011) are indicated by an asterisk.

### Single-nucleotide polymorphism (SNP)

DNA extractions were done on 96 well plates with the DNeasy blood and tissues kit (QUIAGEN). DNA extracts were diluted to obtain a target concentration of 10-15ng/μL. To identify a set of candidate diagnostic SNPs, we used a total of 1380 nuclear ORF contigs/alignments from transcriptomic data for *L. vulgaris* and *L. montandoni* (Stuglik & Babik 2016). Each of these 1380 alignments encompassed the same 13 individuals: six *L. vulgaris* from Romania and Poland), six *L. montandoni* from Romania and Poland) and one *L. helveticus* from France (cf. Supplementary Table 1).

For population-level genotyping, we selected the MassARRAYrray System approach (Sequenom). The MassArray assay consists of an initial locus-specific PCR reaction, followed by single base extension using mass-modified dideoxynucleotide terminators of an oligonucleotide primer which anneals immediately upstream of the polymorphic site of interest. Using MALDI-TOF mass spectrometry, the distinct mass of the extended primer identifies the SNP allele. This technology can genotype a SNP only if it presents biallelic variability and conserved upstream and downstream sequences of about 100 bp each (cf. Gabriel & Ziaugra (2004) and Gabriel et al. (2009) for more details about the Sequenom MassARRAY technology). Among the 1380 alignments examined, only 127 presented sequences meeting the above mentioned specifications and were used to develop 127 multiplexable candidate probes that were preliminarily tested on 20 *L. vulgaris* and 20 *L. helveticus* from different allopatric localities, in Poland (Krakow), Hungary (Pilisz mount and Budapest), Romania (Apuseni), Norway and Sweden for *L. vulgaris*, and Spain (Escobedo, Fresnedo and Poza de la Sal) and the southern part of France (Dordogne, Cazevielle, Ferrières-les-Verries, Notre-Dame-de-Londres and Aumelas) for *L. helveticus*. This screening phase enabled the extraction of 39 SNPs presenting a low amount of missing data and unambiguously distinguishing *L. vulgaris* from *L. helveticus* (cf. Supplementary Table 2 for the sequences of these probes). The selected alignments were used to design a 39-plex array to genotype the 1035 samples. The genotyping was subcontracted to the Genome Transcriptome facility at the Center for Functional Genomics in Bordeaux (CGFB), France.

### Population structure

Sample allocation and admixture level based on the 39 SNPs dataset were assessed with STRUCTURE v2.2 (Pritchard et al. 2000, Falush et al 2003) under models assuming two populations (K=2). STRUCTURE has been shown to be less sensitive to the proportion of hybrids included in the sample, while NEWHYBRIDS (another widely used population genetic programs to address questions related to genetic structure, admixture, and hybridization, Anderson & Thompson 2002) seems to perform slightly better when individuals from both backcross and F_1_ hybrid classes are present in the sample (Vähä & Primmer, 2006). STRUCTURE assigns individuals probabilistically to clusters based on their multilocus genotype. We estimated posterior distributions based on 3 million MCMC generations of which 50% were discarded as burnin. We used a model that considers the possibility of mixed population ancestry and of correlated allele frequencies among populations due to migration or shared ancestry (Falush et al 2003), with an alpha value of 1/k (0.5) (Wang 2017). For comparison, the same analyses were repeated on the microsatellite dataset generated by Johanet et al. (2011, six loci: Tv3ca9, Th09, Tv3Ca19, Th14, Th27, Thca14), to compare membership coefficients of the 78 specimens for which both SNP and microsatellite data were generated. We used the same options for the microsatellite dataset as with the SNPs.

## Results

Samples with three or more missing SNP loci (70 samples) were discarded from the dataset, leading to a final biallelic SNP data-set for 965 usable individual newts, with only 5.3% of missing data (4,256 undetermined SNP genotypes out of a total of 80,730). From the 39 selected SNPs, two showed shared alleles at significant frequencies: SNP 10797 and SNP 10913. For both loci, the “vulgaris” allele was found in *Lissotriton helveticus* in low frequencies that were similar between syntopic (3.3% for SNP 10797 and 13.2% for SNP 10913) and allopatric populations (4.0% and 20.0%, respectively); these two loci are thus not fully diagnostic. In spite of this, the SNPs allowed specific discrimination of every sample, including newly collected as well as previously studied by Johanet et al (2011), and no significant sign of introgression between species was detected, i.e., species-specific alleles of one species were not found in the other. Only two samples collected in two syntopic localities in France revealed an F1-hybrid genotype: one (sample T9-12-LH8f-2, from Roissy-en-Brie) is heterozygous (*vh*) for 37 SNP loci (homozygous for the two remaining SNP loci, one of *vv* type and one *hh* type) and a second sample (T36-14-24-1, from Ecordal) is heterozygous for all the 38 SNP loci available (genotype at one locus missing). No other signs of hybridization or introgression were detected (Fig. 2A and Table 1).

**Fig. 2.**
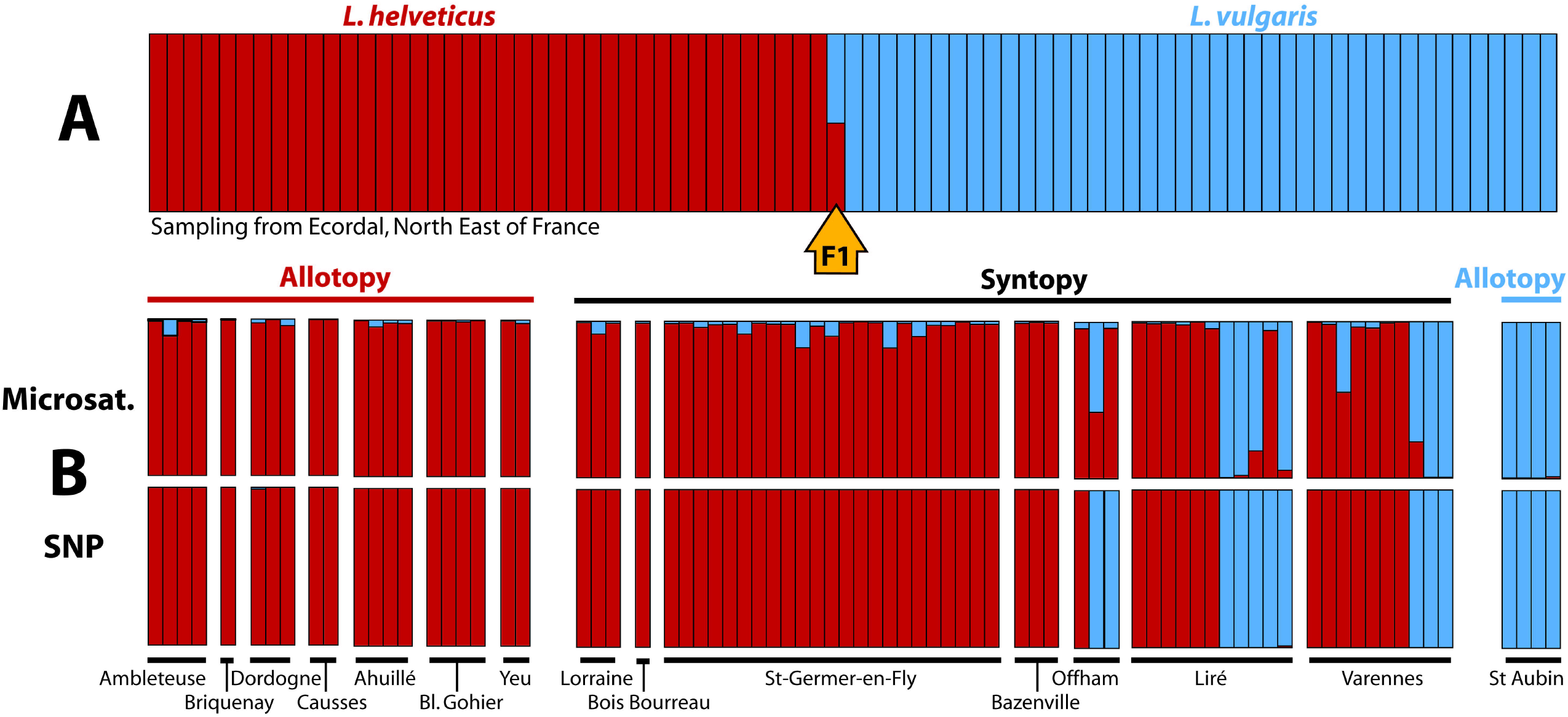
Overview of the genetic structure using STRUCTURE (with K=2). (A) All the samples genotyped with SNPs are unambiguously assigned to one species or another (probability ≥ 0.99), except for two putative F1 hybrid specimens assigned with a probability of 0.50 to both species. Only results from Ecordal (NE France) where one of the two hybrids was detected, are illustrated (cf. Table 1 for overall results). (B) Two alternative Bayesian clustering analyses based on the same set of samples from various localities illustrating the differences between microsatellite and SNP genotyping: the structure inferred from the microsatellite dataset suggests significant introgressive hybridization between both species (above, raw data from Johanet et al. 2011), whereas on the contrary, the structure inferred from the newly generated SNP dataset does not show any signs of admixture between species (below).

In contrast with the SNP-based results, the sample allocation and admixture level presently inferred from the microsatellite dataset previously generated by Johannet et al (2011) suggest a substantial admixture of genes from the other species (i.e., ≥ 0.1) for 11 of the 78 samples presently reanalyzed with SNPs, in accordance with the results of the original publication (See Figure 2B and Supplementary Table 3 for a comparison, for the same 78 samples, of the admixture levels inferred from microsatellites and SNP, respectively).

## Discussion

These results confirm the conclusions of Arntzen et al (1998): *Lissotriton helveticus* and *L. vulgaris* can hybridize in nature but this phenomenon remains rare, as we found only two F1 individuals out of a total of 829 adults from syntopic populations. Such hybrids are likely sterile as we detected no signs of introgression between both species. In addition, SNP analyses of 78 of the samples previously analysed by Johanet et al. (2011), including samples identified as admixed individuals by microsatellite genotypes, detected no sign of hybridization either, suggesting that detection of hybridization and introgression with microsatellites produced biased results that overestimated the frequency of hybridization between these two species (Fig. 2B). The lack of introgression is consistent with the maintenance of a broad sympatric zone and supports the hypothesis that convergence of certain color traits (dorsum, tail) observed in sympatry would primarily be the result of plastic or adaptive responses to environmental variables and not introgression (de Solan et al. 2022). This result also echoes the findings of Drillon et al. (2019) who found hardly any hybridization and no introgression between *Hyla* tree frog species that diverged around 20 Mya. Yet, the possibility of introgressive hybridization can persist for a very long time in newts, as exemplified by *Triturus cristatus* and *T. marmoratus* that hybridize frequently and can still exchange genes in spite of their divergence estimated at around 24 Mya (Wielstra et al. 2011, Arntzen et al. 2021, comparable with the timing of divergence between *L. vulgaris* and *L. helveticus* presented in introduction).

An alternative explanation would be that the SNP genotyping underestimated hybridization and introgression, whereas the microsatellites returned the correct pattern of admixture. Our SNP loci have been selected to be (near) perfectly diagnostic (fixed alternative alleles in both species) and might therefore over-represent genomic regions highly differentiated between species, and thus, indirectly, genes involved in barriers to gene flow (which, by counter-selection, would be less likely to introgress in the long term in other species). On the contrary, microsatellites have been designed and selected independently of their level of divergence between species and should thus not overrepresent genomic regions that resist introgression. This hypothesis could be valid if the microsatellite results were able to detect long-term (historical) interspecific gene flow, as this would indeed be reduced in genomic regions involving barrier loci. However it does not apply to the detection of recent admixture (F1 hybrids and backcrosses), in which case the SNP results are expected to be more robust than the microsatellite results due to a larger number of loci (and unambiguous allelic assignment) in the former. Thus, this hypothesis does not seem able to explain the low frequency of F1s and the total absence of F2 or backcrosses in the SNP dataset, in contradiction with the previous study based on microsatellites.

It thus seems more likely that the microsatellite analyses by Johanet et al. (2011) led to an overestimation of introgression. In general, SNPs have recurrently been shown to outperform microsatellites (e.g. Camacho-Sanchez et al. 2020, Bradbury et al. 2015, Hoffman et al. 2014, Lemopoulos et al. 2019, Sunde et al. 2020, Zimmerman et al. 2020, Szatmári et al. 2021), although it should be emphasized that these works most often involved a far greater number of SNP loci than in the present study (i.e. hundreds or thousands of SNPs *versus* 39 SNPs). More specifically, several recent studies comparing relative performance of SNPs and microsatellites for hybridization detection effectiveness have shown that many hybrids indicated by microsatellites are not validated by SNPs (Daïnou et al. 2017, Szatmári et al, 2020). Assessment of hybridization usually rests, as in the present case, on statistical estimation of the likelihood to observe a given multilocus genotype in a population (assignment methods such as STRUCTURE or NEWHYBRIDS) or comparisons of multivariate axes scores (Johannet at al. 2011, present study). These assignment methods can produce unreliable results when a small number of microsatellites are used or when there is too much missing data (Grünwald et al 2017, Hodel et al. 2017, Yi & Latch 2022). Analyses of datasets simulated under scenarios with different levels of genetic divergence and varying number of loci have for instance shown that two popular model-based Bayesian methods for detecting hybridization (i.e. STRUCTURE and NEWHYBRIDS) require a significantly higher number of loci than commonly applied in microsatellite-based studies (Vähä & Primmer 2006). According to this study, efficient detection of F_1_ hybrid individuals would require for instance the use of 12 or 24 loci (for pairwise *F*_ST_ between hybridizing parental populations of 0.21 or 0.12, respectively), and separating backcrosses from pure individuals would require an even more significant genotyping effort (≥ 48 loci, even when divergence between parental populations are high). In addition to the frequently low number of loci usually involved in microsatellites studies (n = 6 in Johannet et al. 2011), it should be noted that these markers are also notoriously difficult to use to assess hybridization because they are highly polymorphic (from 12 to 35 alleles per locus, with an average of 25 alleles in both species in Johanet et al. 2011) which, coupled with homoplasy, often result in a lack of diagnostic loci (Putman & Carbone 2014, Daïnou et al. 2017, Šarhanová, et al. 2018). Additionally, the study by Johanet et al. (2011) was affected by a significant amount of microsatellite missing data (22.2% for the original dataset). Within the 1924 samples involved, 41.8% were missing at least one third of the data (i.e., 2 or 3 loci out of a total of six) and 11.4% were missing half of the data (i.e., 3 out of 6). When focusing on those samples reanalyzed with SNPs, for which microsatellites suggested substantial admixture of genes from the other species (i.e., ≥ 0.1, n=11), examination of the raw data revealed that all of them had at least one missing locus, with an overall mean amount of missing data reaching 34.8%.

Taken together, these considerations prompt us to conclude that the most parsimonious explanation for the discrepancy between Johanet et al. (2011) on the one hand, and Arntzen et al. (1998) and the present study on the other hand, is the poor performance of NEWHYBRIDS (microsatellite analysis by Johanet et al. 2011) and STRUCTURE (reanalysis of the same dataset in the present study) to reliably assess ancestry on the basis of an inadequate dataset, characterized both by a small number of hypervariable microsatellite loci and a substantial amount of missing data.

Over the past decades, microsatellite genotyping has been the approach favored by the research community to look for hybridization between non-model species. The present study suggests that many of these works may have suffered from biases like those encountered by Johanet et al (2011), and may have overestimated the frequency of this phenomenon, as was the case here. The conclusions from these studies, and the subsequent works based on them, should therefore be carefully reconsidered in the light of their respective datasets coverage. By providing a broad coverage of loci, including diagnostic loci, genomic approaches (such as the SNP approach of the present study) would certainly represent preferable alternatives to microsatellite approaches to investigate introgressive hybridization, and should contribute to refining our understanding about the extent of this phenomenon in nature.

## Acknowledgments

We are grateful to Erwan Guichoux (CGFB) and Patricia Sorouille (CEFE) for their technical support.

## Author contribution statement

P.A.C. and J.S. designed the research concept and realized most of the field work, A.M. performed data extraction and analyses. A.M. and P.A.C wrote the manuscript with contributions to all authors. All authors have read and validated the final version of the manuscript.

## Data archiving

The SNP raw dataset (newly generated) and the microsatellites raw dataset (by Johanet et al. 2011) that support the findings of this study are available in [repository name, DOI(s)] [Cf. XLS files in attachment. To be deposited upon manuscript acceptance].

## Supplementary Information

Supplementary Table 1. List of samples used to identify a set of candidate diagnostic SNPs.

Supplementary Table 2. List and sequences of the 39 DNA probes used for SNP genotyping.

Supplementary Table 3. Assignment coefficients obtained by STRUCTURE for the 78 specimens of *Lissotriton helveticus* (*Lh*) and *L. vulgaris* (*Lv*) living in sympatry and for which both microsatellites and SNP analyses could be performed.

## Bibliography

Adavoudi R, Pilot M (2022) Consequences of Hybridization in Mammals: A Systematic Review. Genes 13:50. https://doi.org/10.3390/genes13010050

Anderson E, Thompson E (2002) A model-based method for identifying species hybrids using multilocus genetic data. Genetics 160:1217–1229. https://doi.org/10.1093/genetics/160.3.1217.

Arntzen JW, Beukema W, Galis F, Ivanović A (2015) Vertebral number is highly evolvable in salamanders and newts (family Salamandridae) and variably associated with climatic parameters. Contrib Zool 84:85–113. https://doi.org/10.1163/18759866-08402001

Arntzen JW, de Wijer P, Jehle R, Smit E, Smit J (1998) Rare hybridization and introgression in smooth and palmate newts (Salamandridae: Triturus vulgaris and T. helveticus). J Zoolog Syst Evol 36:111–122. https://doi.org/10.1111/j.1439-0469.1998.tb00830.x

Arntzen JW, Jehle R, Bardakci F, Burke T, Wallis GP (2009) Asymmetric viability of reciprocal cross hybrids between crested and marbled newts (Triturus cristatus and T. marmoratus). Evolution 63:1191–1202. https://doi.org/10.1111/j.1558-5646.2009.00611.x

Arntzen JW, Jehle R, Wielstra B (2021) Genetic and morphological data demonstrate hybridization and backcrossing in a pair of salamanders at the far end of the speciation continuum. Evol Appl 14:2784–2793. https://doi.org/10.1111/eva.13312

Babik W, Rafiński J (2004) Relationship between morphometric and genetic variation in pure and hybrid populations of the smooth and Montandon’s newt (Triturus vulgaris and T. montandoni). J Zool 262:135–143. https://doi.org/10.1017/S0952836903004369

Babik W, Szymura JM, Rafiński J (2003) Nuclear markers, mitochondrial DNA and male secondary sexual traits variation in a newt hybrid zone (Triturus vulgaris × T. montandoni). Mol Ecol 12:1913–1930. https://doi.org/10.1046/j.1365-294x.2003.01880.x.

Babik W, Branicki W, Crnobrnja-Isailović J, Cogălniceanu D, Sas I, Olgun K, Poyarkov NA, Garcia-París M, Arntzen JW (2005) Phylogeography of two European newt species discordance between mtDNA and morphology. Mol Ecol 14:2475–2491. https://doi.org/10.1111/j.1365-294X.2005.02605.x.

Barraclough TG, Humphreys AM (2015) The evolutionary reality of species and higher taxa in plants: a survey of post-modern opinion and evidence. New Phytol 207:291–296. https://doi.org/10.1111/nph.13232

Barton NH (2001) The role of hybridization in evolution. Mol Ecol 10:551–568. https://doi.org/10.1046/j.1365-294x.2001.01216.x

Beebee TJC, Rowe G, Arntzen JW (1999) Identification of newts (Triturus sp.) and their hybrids using molecular methods. J Zool 249:43–47. doi: 10.1111/j.1469-7998.1999.tb01058.x

Böhme M (2010) Ectothermic vertebrates (Actinopterygii, Allocaudata, Urodela, Anura, Crocodylia, Squamata) from the Miocene of Sandelzhausen (Germany, Bavaria) and their implications for environment reconstruction and palaeoclimate. Palaontol Z 84:3–41. https://doi.org/10.1007/s12542-010-0050-4

Bradbury IR, Hamilton LC, Dempson B, Robertson MJ, Bourret V, Bernatchez L, Verspoor E (2015) Transatlantic secondary contact in Atlantic Salmon, comparing microsatellites, a single nucleotide polymorphism array and restriction-site associated DNA sequencing for the resolution of complex spatial structure. Mol Ecol 24:5130–5144. https://doi.org/10.1111/mec.13395

Camacho-Sanchez M, Velo-Antón G, Hanson JO, Veríssimo A, Martínez-Solano Í, Marques A, Moritz C, Carvalho SB (2020) Comparative assessment of range-wide patterns of genetic diversity and structure with SNPs and microsatellites: A case study with Iberian amphibians. Ecol Evol 10:10353–10363. https://doi.org/10.1002/ece3.6670

Chan WY, Hoffmann AA, van Oppen MJH (2019) Hybridization as a conservation management tool. Conserv Lett 12:e12652. https://doi.org/10.1111/conl.12652

Cogălniceanu D, Stănescu F, Arntzen JW (2020) Testing the hybrid superiority hypothesis in crested and marbled newts. J Zool Syst Evol Res 58:275–283. https://doi.org/10.1111/jzs.12322

Daïnou K., Flot JF, Degen B, Blanc-Jolivet C, Doucet JL, Lassois L, Hardy OJ (2017) DNA taxonomy in the timber genus Milicia: evidence of unidirectional introgression in the West African contact zone. Tree Genet Genomes 13: 90. https://doi.org/10.1007/s11295-017-1174-4

de Solan T, Théry M, Picard D, Crochet P-A, David P, Secondi J (2022) A lot of convergence, a bit of divergence: Environment and interspecific interactions shape body colour patterns in Lissotriton newts. J Evol Biol 35:575–588. https://doi.org/10.1111/jeb.13985

Drillon O, Dufresnes G, Perrin N, Crochet P-A, Dufresnes C. (2019) Reaching the edge of the speciation continuum: hybridization between three sympatric species of Hyla tree frogs, Biol J Linn Soc 126:743–750. https://doi.org/10.1093/biolinnean/bly198

Dufresnes C, Brelsford A, Jeffries D, Mazepa G, Suchan T, Canestrelli D, Nicieza A, Fumagalli L, Dubey S, Martínez-Solano I, Litvinchuk S, Vences M, Perrin N, Crochet P-A. (2021) Mass of genes rather than master genes underlie the genomic architecture of amphibian speciation. PNAS 118:e2103963118. https://doi.org/10.1073/pnas.2103963118

Dudek K, Gaczorek TS, Zieliński P, Babik W (2019) Massive introgression of major histocompatibility complex genes in newt hybrid zones. Mol Ecol 28:4798–4810. https://doi.org/10.1111/mec.15254

Falush D, Stephens M, Pritchard JK (2003) Inference of population structure using multilocus genotype data: linked loci and correlated allele frequencies. Genetics 164:1567–1587.

Gabriel S, Ziaugra L (2004) SNP genotyping using Sequenom MassARRAY 7K platform. Curr Protoc Hum Genet 2:2.12. https://doi.org/10.1002/0471142905.hg0212s42.

Gabriel S, Ziaugra L, Tabbaa D (2009) SNP genotyping using the Sequenom MassARRAY iPLEX platform. Curr Protoc Hum Genet.2:2.12. https://doi.org/10.1002/0471142905.hg0212s60.

Gherghel I, Strugariu A, Ambrosă I-M, Zamfirescu □ (2012) Updated distribution of hybrids between Lissotriton vulgaris and Lissotriton montandoni (Amphibia: Caudata: Salamandridae) in Romania. Acta Herpeto 7: 49–55. https://doi.org/10.13128/Acta_Herpetol-10224

Griffiths RA (1987) Microhabitat and seasonal niche dynamics of smooth and palmate newts, Triturus vulgaris and T. helveticus, at a pond in mid-Wales. J Anim Ecol 56:441–451. https://doi.org/10.2307/5059

Grünwald NJ, Everhart SE, Knaus BJ, Kamvar ZN (2017) Best Practices for Population Genetic Analyses. Phytopathology 107:1000–1010. https://doi.org/10.1094/PHYTO-12-16-0425-RVW

Harrison RG, Larson EL (2014) Hybridization, Introgression, and the Nature of Species Boundaries. J Hered 105:795–809, https://doi.org/10.1093/jhered/esu033

Hirashiki C, Kareiva P, Marvier M (2021) Concern over hybridization risks should not preclude conservation interventions. Conserv Sci Pract 3:e424. https://doi.org/10.1111/csp2.424

Hodel RGJ, Chen S, Payton AC, McDaniel SF, Soltis P, Soltis DE (2017) Adding loci improves phylogeographic resolution in red mangroves despite increased missing data: comparing microsatellites and RAD-Seq and investigating loci filtering. Sci Rep 7:17598. https://doi.org/10.1038/s41598-017-16810-7

Hoffman JI, Simpson F, David P, Rijks JM, Kuiken T, Thorne MAS, Lacy RC, Dasmahapatra KK (2014) High-throughput sequencing reveals inbreeding depression in a natural population. PNAS 111:3775. https://doi.org/10.1073/pnas.1318945111

Jehle R, Arntzen JW, Burke T, Krupa AP, Hödl W (2001). The annual number of breeding adults and the effective population size of syntopic newts (Triturus cristatus, T. marmoratus). Mol Ecol 10:839–850. https://doi.org/10.1046/j.1365-294X.2001.01237.x

Johanet A, Secondi J, Lemaire C (2011) Widespread introgression does not leak into allotopy in a broad sympatric zone. Heredity 106:962–972. https://doi.org/10.1038/hdy.2010.144

Kim D, Bauer BH, Near TJ (2022) Introgression and Species Delimitation in the Longear Sunfish Lepomis megalotis (Teleostei: Percomorpha: Centrarchidae). Syst Biol 71:273–285, https://doi.org/10.1093/sysbio/syab029

Lemopoulos A, Prokkola JM, Uusi-Heikkilä S, Vasemägi A, Huusko A, Hyvärinen P, Koljonen M-L, Koskiniemi J, Vainikka A (2019) Comparing RADseq and microsatellites for estimating genetic diversity and relatedness — Implications for brown trout conservation. Ecol Evol 9:2106–2120. https://doi.org/10.1002/ece3.4905

Mallet J (2005) Hybridization as an invasion of the genome. Trends Ecology Evol 20:229–237. https://doi.org/10.1016/j.tree.2005.02.010.

Mayr E (1963) Animal Species and Evolution. Belknap; Cambridge, MA, USA: 1963.

McFarlane SE, Pemberton JM (2019) Detecting the True Extent of Introgression during Anthropogenic Hybridization. Trends Ecol Evol 34:315–326. https://.doi.org/10.1016/j.tree.2018.12.013.

Moran BM, Payne C, Langdon Q, Powell DL, Brandvain Y, Schumer M (2021) The genomic consequences of hybridization. eLife 10:e69016. https://doi.org/10.7554/eLife.69016

Niedzicka ME, Głowacki BM, Zieliński P, Babik W (2020). Morphology is a poor predictor of interspecific admixture – the case of two naturally hybridizing newts Lissotriton montandoni and Lissotriton vulgaris (Caudata: Salamandridae). Amphib Reptil 41:489–500. https://doi.org/10.1163/15685381-bja10019

Niedzicka M, Dudek K, Fijarczyk A, Zieliński P, Babik W (2017). Linkage Map of Lissotriton Newts Provides Insight into the Genetic Basis of Reproductive Isolation. G3: Genes Genomes Genet 7:2115–2124. https://doi.org/10.1534/g3.117.041178.

Pabijan M, Zieliński P, Dudek K, Stuglik M, Babik W (2017). Isolation and gene flow in a speciation continuum in newts. Mol Phylogenet Evol 116:1–12. https://doi.org/10.1016/j.ympev.2017.08.003

Pfennig KS, Kelly AL, Pierce AA (2016) Hybridization as a facilitator of species range expansion. Proc R Soc B 283:20161329–9. https://doi.org/10.1098/rspb.2016.1329

Pritchard JK, Stephens M, Donnelly P (2000) Inference of population structure using multilocus genotype data. Genetics 155:945–959. https://doi:10.1093/genetics/155.2.945

Putman AI, Carbone I (2014) Challenges in analysis and interpretation of microsatellite data for population genetic studies. Ecol Evol 22:4399–4428. https://doi:10.1002/ece3.1305.

Rage JC & Bailon S (2005) Amphibians and squamate reptiles from the late early Miocene (MN 4) of Béon 1 (Montréal-du-Gers, southwestern France). Geodiversitas 27:413–441.

Šarhanová P, Pfanzelt S, Brandt R, Himmelbach A, Blattner FR (2018) SSR-seq: Genotyping of microsatellites using next-generation sequencing reveals higher level of polymorphism as compared to traditional fragment size scoring. Ecol Evol 8:10817–10833. https://doi.org/10.1002/ece3.4533

Schlüpmann M, Weber G, Lipscher E, Veith M (1999) Nachweis eines freinlandbastardes von Teichmolch (Triturus vulgaris) und Fadenmolch (Triturus helveticus). Z Feldherpetol 6:203–217.

Seabra, SG, Silva, SE, Nunes, VL, Sousa VC, Martins J, Marabuto E, Rodrigues ASB, Pina-Martins F, Laurentino TG, Rebelo MT, Figueiredo E, Paulo OS (2019) Genomic signatures of introgression between commercial and native bumblebees, Bombus terrestris, in western Iberian Peninsula—Implications for conservation and trade regulation. Evol Appl 12:679–691. https://doi.org/10.1111/eva.12732

Secondi J, Johanet A, Pays O, Cazimajou F, Djalout Z, Lemaire C (2010) Olfactory and visual species recognition in newts and their role in hybridization. Behaviour 147:1693–1712.

Sillero N, Campos J, Bonardi A, Corti C, Creemers R, Crochet PA, Crnobrnja Isailović J, Denoël M, Ficetola GF, Gonçalves J, Kuzmin S, Lymberakis P, de Pous P, Rodríguez A, Sindaco R, Speybroeck J, Toxopeus B, Vieites DR, Vences M (2014) Updated distribution and biogeography of amphibians and reptiles of Europe. Amphib Reptil 35:1–31. https://doi.org/10.1163/15685381-00002935

Sunde J, Yildirim Y, Tibblin P, Forsman A (2020) Comparing the performance of microsatellites and RADseq in population genetic studies: Analysis of data for pike (Esox lucius) and a synthesis of previous studies. Front Genet 11:218. https://doi.org/10.3389/fgene.2020.00218

Steensels J, Gallone B, Verstrepen KJ (2021) Interspecific hybridization as a driver of fungal evolution and adaptation. Nat Rev Microbiol 19:485–500. https://doi.org/10.1038/s41579-021-00537-4

Steinfartz S, Vicario S, Arntzen JW, Caccone A (2007) A Bayesian approach on molecules and behavior: reconsidering phylogenetic and evolutionary patterns of the Salamandridae with emphasis on Triturus newts. J Exp Zool B Mol Dev Evol. 2308:139–62. https://doi.org/10.1002/jez.b.21119.

Stuglik MT, Babik W (2016) Genomic heterogeneity of historical gene flow between two species of newts inferred from transcriptome data. Ecol Evol 6:4513–4525, https://doi.org/10.1002/ece3.2152

Szatmári L, Cserkész T, Laczkó L, Lanszki J, Pertoldi C, Abramov AV, Elmeros M, Ottlecz B, Hegyeli Z, Sramkó G (2021) A comparison of microsatellites and genome-wide SNPs for the detection of admixture brings the first molecular evidence for hybridization between Mustela eversmanii and M. putorius (Mustelidae, Carnivora). Evol Appl 14:2286–2304. https://doi.org/10.1111/eva.13291

Taylor SA, Larson E (2019) Insights from genomes into the evolutionary importance and prevalence of hybridization in nature. Nat Ecol Evol 3:170–177. doi: 10.1038/s41559-018-0777-y.

Taylor EB, Boughmann JW, Greonenboomm M, Sniatynksi D, Schluter D, Gow JL (2006) Speciation in reverse: morphological and genetic evidence of the collapse of a three-spined stickleback Gasterosteus aculeatus) species pair. Mol Ecol 15:343–355.

Vähä JP, Primmer CR (2006) Efficiency of model-based Bayesian methods for detecting hybrid individuals under different hybridization scenarios and with different numbers of loci. Mol Ecol 15:63–72. https://doi.org/10.1111/j.1365-294X.2005.02773.x

Wacker S, Aronsen T, Karlsson S, Ugedal O, Diserud OH, Ulvan EM, Hindar K, Næsje TF (2021) Selection against individuals from genetic introgression of escaped farmed salmon in a natural population of Atlantic salmon. Evol Appl 14:1450–1460. https://doi.org/10.1111/eva.13213

Wang J (2017) The computer program structure for assigning individuals to populations: easy to use but easier to misuse. Mol Ecol Resour 17:981–990. https://doi.org/10.1111/1755-0998.12650

Wielstra B, Arntzen JW (2011) Unraveling the rapid radiation of crested newts (Triturus cristatus superspecies) using complete mitogenomic sequences. BMC Evol Biol 11:162. https://doi.org/10.1186/1471-2148-11-162

Wielstra B, Canestrelli D, Cvijanović M, Denoël M, Fijarczyk A, Jablonski D, Liana M, Naumov B, Olgun K, Pabijan M, Pezzarossa A, Popgeorgiev G, Salvi D, Si Y, Sillero N, Sotiropoulos K, Zieliński P, Babik W (2018) The distributions of the six species constituting the smooth newt species complex (Lissotriton vulgaris sensu lato and L. montandoni)–an addition to the New Atlas of Amphibians and Reptiles of Europe. Amphib Reptil 39:252–259.

Yi X, Latch EK (2022) Nonrandom missing data can bias Principal Component Analysis inference of population genetic structure. Mol Ecol Resour 22:602–611. https://doi.org/10.1111/1755-0998.13498

Zieliński P, Nadachowska-Brzyska K, Wielstra B, Szkotak R, Covaciu-Marcov SD, Cogălniceanu D, Babik W (2013) No evidence for nuclear introgression despite complete mtDNA replacement in the Carpathian newt (Lissotriton montandoni). Mol Ecol 22:1884–1903.

Zieliński P, Stuglik MT, Dudek K, Konczal M, Babik W (2014) Development, validation and high-throughput analysis of sequence markers in nonmodel species. Mol.Ecol. Resour. 14, 352–360.

Zieliński P, Nadachowska-Brzyska K, Dudek K, Babik W (2016) Divergence history of the Carpathian and smooth newts modelled in space and time. Mol. Ecol. 25, 3912–3928.

Zimmerman SJ, Aldridge CL, Oyler-McCance SJ (2020) An empirical comparison of population genetic analyses using microsatellite and SNP data for a species of conservation concern. BMC Genomics 21, 382. https://doi.org/10.1186/s12864-020-06783-9

